# HIF-1α stabilization modified by glutaredoxin-1 is critical for intestinal angiogenesis in NEC pathogenesis

**DOI:** 10.1101/2021.08.30.458202

**Authors:** Yunfei Zhang, Xiao Zhang, Bing Tian, Xionghui Ding, Cuilian Ye, Chunbao Guo

**Affiliations:** Department of General and Neonatal Surgery, Children’s Hospital of Chongqing Medical University, Chongqing, China; Department of Burn, Children’s Hospital of Chongqing Medical University, Chongqing, ChinaS; School of Pharmacy and Bioengineering, Chongqing University of Technology, Chongqing, 400054, P.R. China; Ministry of Education Key Laboratory of Child Development and Disorders, Children’s Hospital of Chongqing Medical University, Chongqing, China

**Keywords:** necrotizing enterocolitis, Hypoxia inducible factor, glutaredoxin-1, vascular endothelial growth factor, intestinal microvascular

## Abstract

**Background:** Hypoxia inducible factor (HIF-1α) are essential in the pathogenesis of necrotizing enterocolitis (NEC), which is stabilized by Grx1 deletion. Until now, the mechanism of HIF-1α in the intestinal microcirculation in NEC is not well defined. We intend to investigate the role of HIF-1α in the development of NEC in regulating the microcirculation and the following vasodilatory signal, VEGF.

**Materials and methods:** Experimental NEC was induced in full-term C57BL/6 mouse and Grx1^-/-^ pups through the formula gavage and hypoxia technique. The HIF-1α signal was blocked utilizing the HIF-1α inhibitor, YC-1. Intestinal tissues were collected at predetermined time points for the assessment of intestinal microcirculation and the HIF-1α activity involved signal.

**Results:** We found that NEC inducement impaired the intestinal microcirculation, but intestinal blood flow and capillary density were ameliorated in Grx1^-/-^ mice, which was associated with the GSH-protein adducts of HIF-1α in the intestinal tissue. Grx1 ablation could also promote vascular endothelial growth factor (VEGFA) production in the intestinal tissue. This intestinal microvascular improvement was not found in the HIF-1α inhibited mice, suggesting the HIF-1α dependent manner for intestinal microcirculatory perfusion.

**Conclusion:** The current data demonstrated that HIF-1α signaling is involved in the intestinal microvascular modification during the pathogenesis of NEC, suggesting that targeting with HIF-1α might be a promising strategy for NEC treatment.

## Introduction

Necrotizing enterocolitis (NEC) is the common and serious devastating gastrointestinal disease in premature infants[1], with common pathological changes of the intestine inflammation, ischemia and necrosis[2]. The etiology of NEC is not fully addressed and so the management of NEC remains challenging. Many highly risk factors suggest that oxygen transport to intestinal cells plays a critical role in NEC, include Birth asphyxia, congenital heart disease, blood transfusion, maternal preeclampsia, intrauterine growth restriction[3, 4]. Hypoxia caused by transient ischemic injury leading to multitude of inflammatory responses in the gut is thought to be a critical initial insult[2].

Hypoxia-inducible factor HIF-1α, a transcription factor activated by Hypoxia, is essential in regulating hypoxia-induced gene expression that triggers various cellular, systemic, and metabolic responses necessary for tissues to adapt to low oxygen conditions[5]. HIF-1α contributes to O_2_ homeostasis, wound healing, erythropoiesis, cell survival, and regulation of vascular tone, which is a heterodimer consisting of constitutively expressed α and β subunits[6]. The exact signaling mechanisms of HIF-1α in NEC are not fully understood.

Post-translational modification critically determines HIF-1α activity and stability. The degradation of HIF-1α is mediated by prolyl hydroxylase domain enzymes (PHDs) that hydroxylate HIF-α subunit on two proline residues. Hydroxylation serves as a recognition motif for the von Hippel-Lindau tumor suppressor protein, which recruits an E3 ubiquitin ligase complex leading to rapid proteasomal degradation of HIF-α subunit[7]. In hypoxia, HIF-1α heterodimerization with HIF-1β leading to form heterodimer and translocated into the nucleus and binds to HIF response elements, the promoter of HIF target genes, thereby activating transcription of genes that regulate oxygen homeostasis[8].

Oxidants oxidative modifications of redox sensitive cysteine thiols to transduce cellular signaling, the reversible modification by abundant glutathione, called S-glutathionylation (or GSH adducts)[9]. Glutaredoxin-1 (Grx1) is a cytosolic enzyme that specifically catalyzes the *S*-glutathionylated proteins to removal of GSH adducts[10, 11]. GSH adducts preventing interaction with pVHL to stabilize HIF-1α protein in vitro, and GSH adducts on Cys^520^ of HIF-1α be recognized by mass spectrometry analysis. GSH adducts stabilize HIF-1α, and the ischemic revascularization improved by genetic deletion of Grx1 [12]. Grx1 overexpression removed GSH adducts and attenuated revascularization in hind limb ischemia, which may be related to the VEGF signal path[13]. The potential role of GSH-protein adducts in vivo target proteins in NEC has not been elucidated. we hypothesized that the degradation of HIF-1α may be prevented by GSH-protein adducts and that HIF-1α activity might be controlled by Grx1 reversing GSH adducts.

Research reported that using Dimethyloxalylglycine (DMOG) to stabilizing HIF-1α protects against NEC by improving intestinal microvascular integrity and promoting cell proliferation of intestinal endothelial[14]. The expression of pro-angiogenic genes and their corresponding receptors are up regulated by activated HIF-1α, including VEGF and its receptors[15]. The VEGF family members (VEGF-A, -B, -C, -D, and -E) are considered to be one of the most important regulators of angiogenesis and studies have shown that its major angiogenic function is mediated by the interaction between VEGFA and VEGF receptor 2 (VEGFR-2)[16]. Here, we demonstrated that HIF-1α stabilized by GSH adducts as a consequence of depletion of Grx1 and protected against NEC by promoting intestinal microvasculature development and intestinal perfusion, which related to HIF-1α/VEGF signaling pathway.

## Materials and Methods

### Animal models and drug treatment

All the experimental protocol was approved by the institute animal care and use committee of Chongqing Medical University. The Grx1^-/-^ mice (C57BL/6J genetic background) were kindly gifted from Prof. Jingyu Li (Sichuan University). The Wild-type (WT) mice (C57BL/6J) were obtained from the experimental animal center of Chongqing Medical University. The six-day-old Grx1-/- mice and WT mice were collected, randomly divided into the control and NEC group separately. For the control group, the pups stayed with the dam without any other treatment. For the NEC group, the mice were subjected to formula gavage containing 15 g of Similac 60/40 (Ross Pediatrics, Columbus, USA) and 75 mL of puppy canine milk replacement (Pet-Ag, Hampshire, USA) at an interval of 4 h, and oral administration of lipopolysaccharide (LPS 5 mg/kg) once per day. The mice underwent hypoxia (100% N2) for 60 s and hypothermia (4□) for 10 min three times per day for 3 days. To study the effect of HIF-1α inhibition on NEC incidence, the HIF-1α inhibitor YC-1 solution (dissolved in 1% dimethyl sulfoxide [DMSO]) were intraperitoneal injected to the litters of newborn pups at a dose of 2 mg/kg body weight daily for 3 days[17, 18]. All the mice were euthanized after modeling, and the distal ileum tissues were collected for experimental detection.

### Human small intestine

Human intestinal tissue specimens were collected from the preterm or full-term infants in the Children’s Hospital of Chongqing Medical University in 2020. The research was conducted with the ethical standards prescribed by the Helsinki Declaration and the protocol was approved by the Institutional Review Board of children’s hospital, Chongqing Medical University. The ileum samples were obtained from infants with acute active NEC, who undergone emergency laparotomy (n = 8) and age-matched infants (n = 6) undergoing surgery for duodenal atresia, ileal atresia, or intestinal reanastomosis.

### Morphological and histological assessment

The formalin-fixed terminal ileum samples were subjected to microscopic examination of H&E stained sections at 4 μm. Histological severity was scored by two investigators blinded to the treatment groups using a standard histologic scoring system reported previously[19].

### Confocal immunofluorescence

The protocol of the immunofluorescence evaluation immunofluorescence for the snap-frozen slides was described previously by our laboratory. Briefly, the sections in different treatment groups were first subjected to antigen retrieval and then incubated in appropriate primary antibodies at 4□ overnight: CD31 Rabbit pAb (383815, zenbio, China), HIF-1 alpha Rabbit pAb (382705, zenbio, China). The sections were followed with incubation with the appropriate secondary antibodies corresponding to the species of the primary antibodies. The nuclear staining was performed using 4’,6-diamidino-2-phenylindole (DAPI, Sigma). Finally, the slides were visualized with a confocal fluorescence microscopy. After surveying the entire section from 6 mice under each experimental condition, 20 high-power fields were examined for each tissue slide. Quantitative analysis of the fluorescence intensity and capillaries in the intestinal was performed for the confocal images using WCIF ImageJ software (National Institutes of Health, Bethesda, MD).

### Western blot analysis

The snap-frozen intestinal tissues were homogenized and centrifuged for the supernatants collection. Following total protein concentration measurement (bicinchoninic acid [BCA] method). Equal amounts of proteins (40 mg) were first loaded on SDS-PAGE gels for electrophoresis and then transferred onto polyvinylidene difluoride (PVDF) membranes (IPFL00010, Millipore, US) for primary antibody incubation. The membranes were blocked with QuickBlock™ Western (Beyotime Institute of Biotechnology, China) and then incubated with primary antibodies at 4□ overnight: vascular endothelial growth factor A (VEGFA) (26157-1-AP, Proteintech), HIF-1 alpha Rabbit pAb (382705, zenbio, China), beta Actin (20536-1-AP, Proteintech, US). The relevant secondary antibodies conjugated with horseradish peroxidase were further incubated at room temperature for 1 h to reveal the immunoreactive bands. The relative intensities of the bands was scanned and archived using a Kodak Scientific Imaging System (Kodak, Rockville, MD), and the optical density was measured utilizing ImageJ software.

### Intestinal epithelial cells (IEC) migration and proliferation

To measure intestinal epithelial cells (IEC) migration and proliferation, the pups with different management were injected intraperitoneally with bromodeoxyuridine (50 mg/kg, 5-BrdU; MCE, US) and sacrificed 18 h later. The terminal ileum samples were subjected to immunofluorescence stain using anti-BrdU antibody. The quantification protocol for IEC migration and proliferation has been described previously[20].

### Statistical analysis

The quantitative data analysis was conducted utilizing GraphPad Prism software (version 4) (GraphPad, San Diego, CA). Continuous parametric data are presented as the mean±SEM, as data were normally distributed. Statistically significant differences were determined using Student’s t-test or ANOVA followed by Bonferroni’s multiple comparison test, as appropriate. P values <0.05 was considered as significance.

## Results

### Intestinal HIF-1a is decreased in human NEC

Previous research has indicated that HIF-1a is required for the angiogenesis, so we speculated that the HIF-1a may be closely related to the microvascular vessels damage and severity of small bowel injury in the NEC pathogenesis.

The location of necrosis and perforation presented with severe damage that shown the lowest or even absent in the levels of HIF-1α protein and VEGFA, a key angiogenic factor among multiple VEGF family members in NEC children intestinal compared with control group (Fig. 1A, 1B, 1C).

**Fig. 1.**
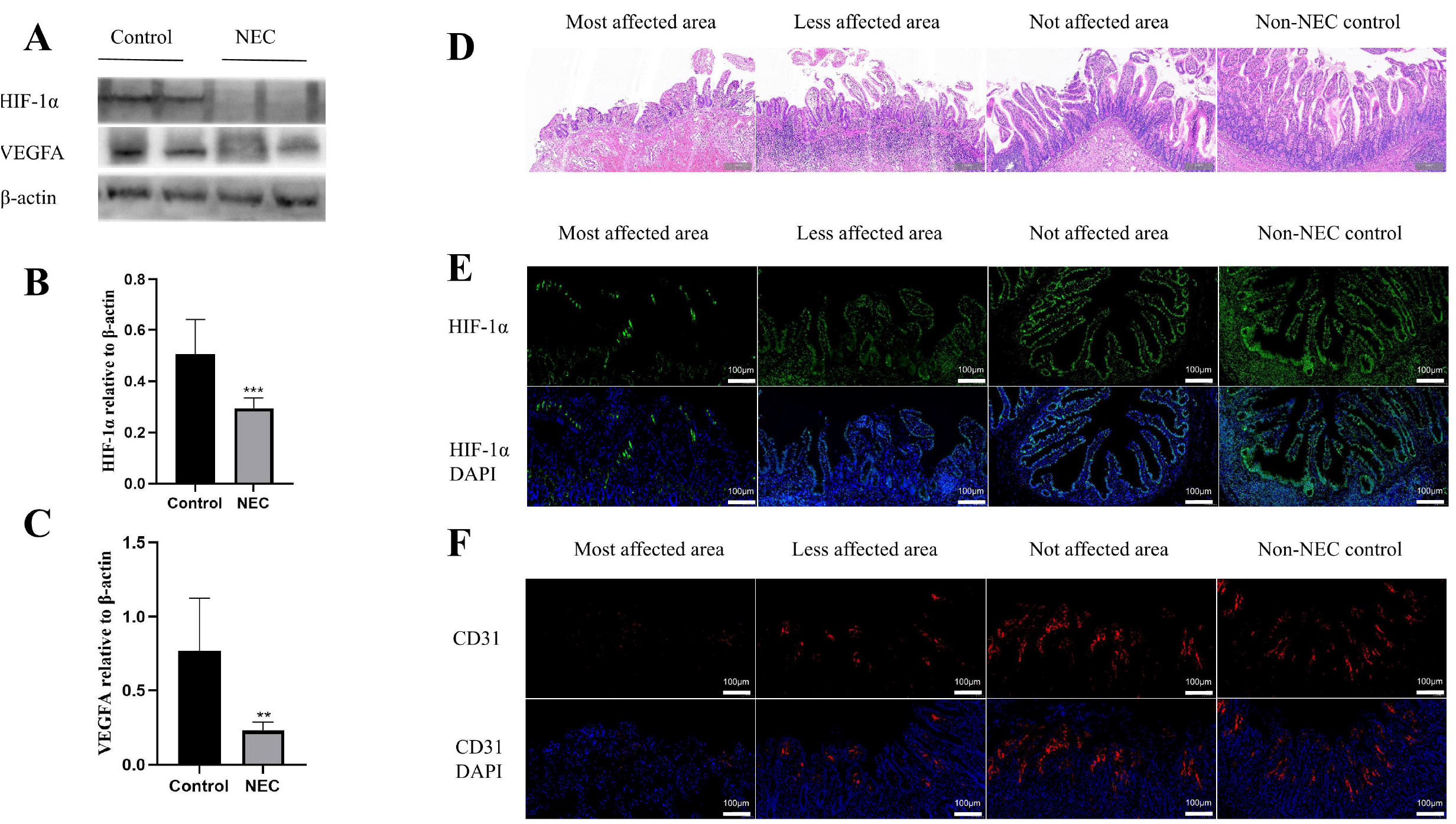
NEC patients are associated with intestinal hypoxia and impairment of microvasculature. A. HIF-1α and VEGFA protein was analyzed in human small intestinal tissues utilizing western blot. B. Band densitometry value of HIF-1α normalized to β-actin are presented. C. Band densitometry value of VEGFA normalized to β-actin are presented. D. Representative morphology of human ileum using H&E staining. We divide intestine into three regions base on the distance to the area of necrosis: the most affected ileum, the less affected ileum and the not affected ileum. Scale bar = 200 μm. Representative intestinal immunofluorescence staining of HIF-1α (E) and CD31 (F) were presented; NEC (n = 8) and non-NEC control (n = 6). Scale bar = 100 μm. **p<0.05, ***p < 0.001 VS Control, unpair T-test.

We next compared the different portions of the terminal ileum at varying distance from the most damaged area resected from human infants with NEC under hematoxylin and eosin staining. In the most affected intestinal area of NEC, the histological feature exhibited most severe injury. The ileum farther away from this most affected area presented with almost normal features resembling characteristic with non-NEC control infants (Fig. 1D).

HIF-1α staining using immunofluorescence measurements was further developed at the most affected terminal ileum., The staining for HIF-1α was noted lowest or even absent expression in the severe damaged intestinal tissues in comparison to the less involved, not affected areas and non-NEC controls (Fig. 1E).

Remarkably, the microvascular vessels and vascular endothelial marker (Platelet endothelial cell adhesion molecule, CD31) were reduced or even absent completely in the damaged villi compared to the less affected ileum, suggesting the severity of small bowel injury (Fig. 1F). The microvascular derangements are maximal in the most affected intestinal area and gradually regains normal features far away from the affected intestinal area (Fig. 1F). These findings suggested that the HIF-1α regulate the formation of blood vessels, that may account for the reduce of endothelial cells and microvascular vessels, presented in the NEC.

### Regulation of HIF-1a stabilization by Grx1 during NEC development

The tripeptide glutathione (GSH) is implicated in NEC pathological mechanism through the redox regulation in combating cellular oxidation (i.e., inflammation). The oxidative stress evaluated by the GSH and GSSG levels in intestinal tissue of experimental NEC mice showed that the GSH level decreased slightly compared with corresponding control mice, but Grx1 ablation significantly decreased the GSH content in NEC mice (Fig. 2B). GSSG content exhibited as the same level in experimental NEC mice compared with corresponding control mice, Grx1 ablation increased the GSSG content significantly in NEC mice (Fig. 2B). Also, in Grx1^-/-^ NEC mice, the GSH:GSSG ratio decreased significantly when exposed to NEC stress regimen (Fig.2A). The protein-GSH adduct level can be cleaved by Grx1, which may therefore constitute an important regulatory switch of HIF-1α stabilization through Grx1 dependent catalysis [16].

**Fig. 2.**
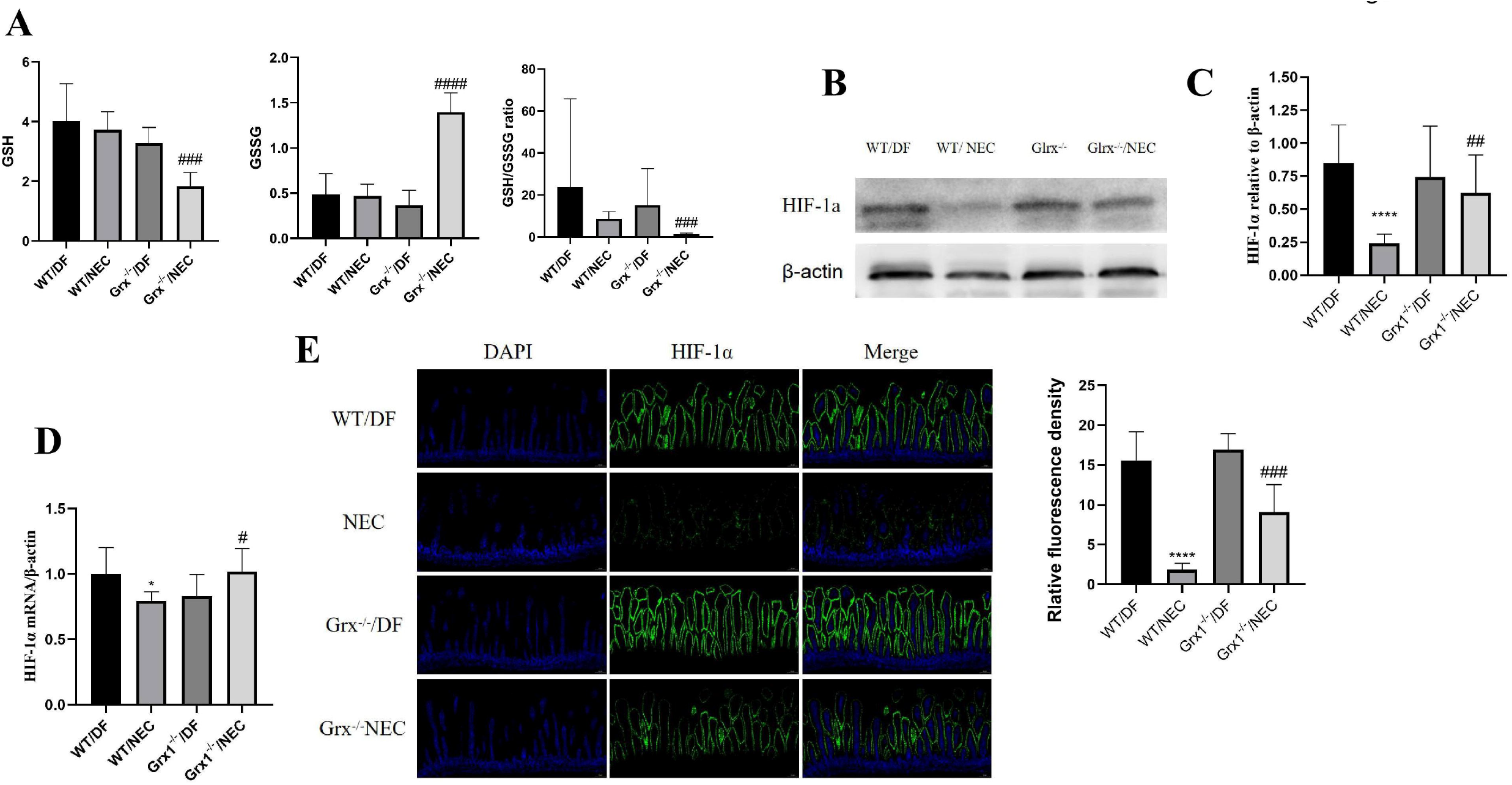
Grx1-depletion was responsible for HIF-1a stabilization through GSH-protein adducts A. The GSH, GSSG and the GSH to GSSG ratio in the intestinal homogenates were determined as described in the instructions of the kit. Independent experiments were repeated three times; Average values are shown; bars, SEM. B. Western blot analyses for HIF-1α from intestinal homogenates of mice. Right panel: The relative intensity of HIF-1a to β-actin. C. HIF-1α mRNA was detected in the intestinal tissues by RT-PCR. D. Representative IF staining of HIF-1α in intestinal tissue of mice treated as indicated. Scale bar = 200 μm. HIF-1α, green; DNA, blue. Right panel: The relative fluorescence density of HIF-1a was quantified. *P < 0.05, ****P < 0.0001 vs WT/DF; ^#^P < 0.05, ^##^P < 0.01, ^###^P < 0.001, ^####^P < 0.0001 vs WT/NEC; one-way ANOVA.

We next assessed the HIF-1α stabilization level utilizing western blot to investigate the effect of GSH adducts removal in Grx1^-/-^ mice. The NEC-inducing regimen apparently down regulated the HIF-1α level (Fig. 2B, 2C). Importantly, the levels of HIF-1α were increased in Grx1^-/-^ pups compared with WT pups (Fig. 2B, 2C), confirming the function of Grx1 in regulatory switch for HIF-1α stabilization. The mRNA levels of HIF-1α did not be affected by Grx1 ablation (Fig. 2D), implying that HIF-1α was regulated by Grx1 at the protein modification level. These results suggested that Grx1 play an important role in controlling the HIF-1α signal pathway through GSH adducts regulation.

To further address the relationship between HIF-1α and NEC, Immunofluorescence staining was performed on serial intestinal sections. The main expression of HIF-1α is located in intestinal epithelial. Relative intensity of fluorescence for HIF-1α reduced remarkable in the NEC mice compared with corresponding control mice; in contrast, HIF-1α expression increased after Grx1 ablation (Fig. 2E).

### Inhibition of HIF-1 signaling abrogates the protective effect of Grx1 ablation on mortality and intestinal injury

We previously found that Grx1 ablation are more resistance to NEC and this phenotype should be involved in the HIF-1 signal. To determine whether HIF-1α signaling molecules is indispensable for Grx1 ablation-induced NEC protection, we next evaluated the roles of HIF-1α in the NEC development.

The typical pathological features of all indicated managements were shown in (Fig. 3A). NEC-inducing regimen caused severe intestinal injury in WT/NEC mouse, shown as villi epithelial sloughing, loss of villi, intestinal necrosis or perforation, but Grx1 ablation mitigated this degree of injury, just presented as disarrangement of villus enterocytes, blunting or edema of villi. The pathological scores was used to assess the degree of intestinal injury. As shown in Fig. 3B, the pathological scores increased after NEC treatment, but Grx1 ablation reduced the pathological scores significantly in NEC group. As expected, Grx1 ablation-mediated protective effect was abolished when HIF-1α inhibitors, YC-1 were given in addition to Grx1 ablation (Fig. 3A, 3B).

**Fig. 3.**
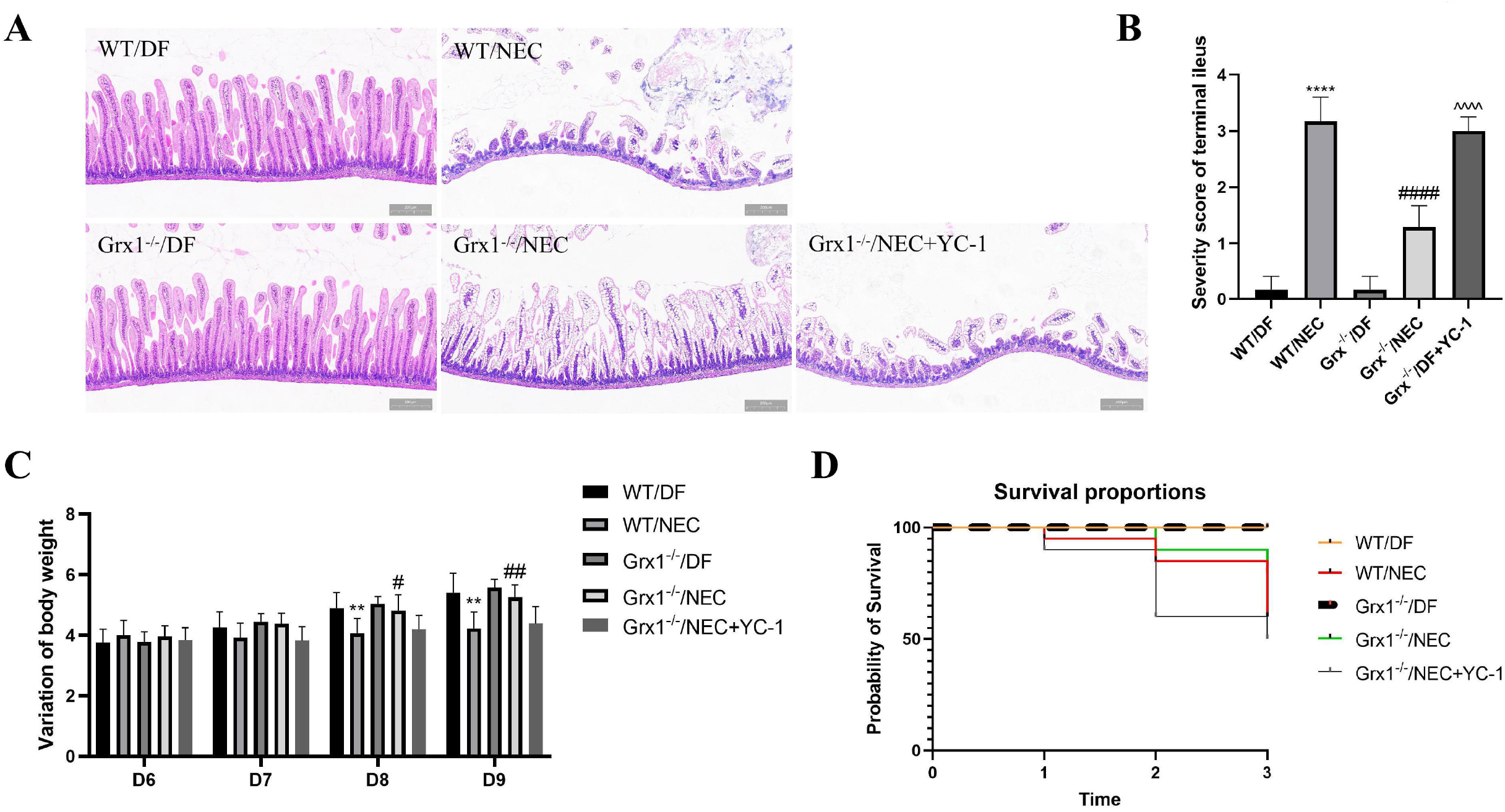
HIF-1a stabilization by Grx1 knockout attenuate the severity of NEC mice A. H&E stanning of intestinal section for morphometric analyses was performed. Representative image of each group is shown. Scale bar = 200 μm. B. Histogram of NEC severity scores from the histopathological evaluation of intestinal (n = 6 animals for each group, 6 fields/animal). C. The body weight change over the three-day protocol in all groups at indicated time point. D. Kaplan-Meier curves for overall survival of each group is shown. The data represent 10-20 mice per group at each time point. Data represent the mean ± SEM. **P < 0.01, ****P < 0.0001 vs WT/DF; ^#^P < 0.05, ^##^P < 0.01, ^####^P < 0.0001 vs WT/NEC; ^^^^P < 0.0001 vs Grx1^-/-^/NEC+YC-1(one-way ANOVA).

Because the intestinal injury is associated with uptake nutrition, we next studied the change of body weight. Dam feed neonatal mice continuously gained weight throughout the entire experiment, NEC treatment reduced weight gain, but this trend was mitigated in Grx1^-/-^ mice, whereas the YC-1 restricted the effect of Grx1 ablation(Fig. 3C). Similarly, the survival curves of the pups indicated that postpartum survival was reduced by NEC treatment, but Grx1 ablation reversed this trend. The YC-1 could abolish the survival benefit of Grx1 ablation in experimental NEC mice (Fig. 3C, 3D). Taken together, our data revealed that the Grx1 ablation counteracted intestinal injury in NEC via HIF-1α signal.

### Stabilize HIF-1α by Grx1 ablation promoted Migration and proliferation, and both effects are abrogated by HIF-1α inhibition

We next investigated whether HIF-1α modulates small intestinal endothelial cell Migration and proliferation, which was involved in NEC pathogenesis. The endothelial cells were tracked within intestinal tissue sections by immunostain for BrdU (Fig. 4A). The normal Intestinal epithelial cell (IEC) migration feature was presented in DF pups (14.81 μm/h) and that was severely impaired in intestine from pups subjected to NEC (4.87 μm/h), whereas Grx1 ablation markedly ameliorated the severely impaired IEC (10.12 μm/h) (Fig. 4C). Consistant with these findings, when IEC migration was calculated as FLE/mucosal thickness ×100%, IEC migration decreased significantly from 80.48% in DF pups to 53.96% in NEC pups and was significantly increased to 68.15% in Grx1-/- pups (Fig. 4D). The IEC proliferation was determined by the BrdU-positive cells, which was also decreased significantly in NEC stressed pups and increased in Grx1 ablation condition (Fig. 4E).

**Fig. 4.**
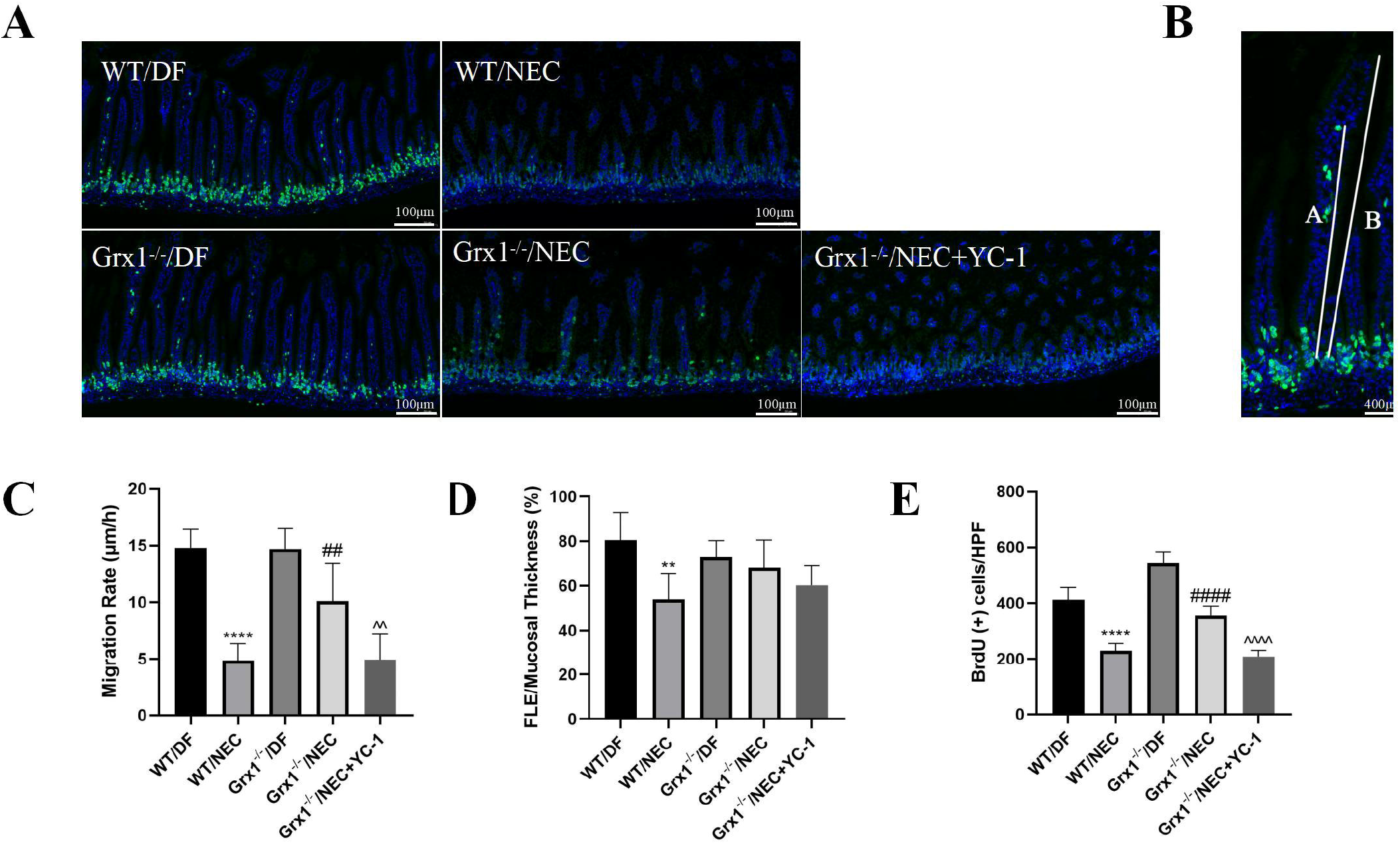
The microvascular features in the experimental NEC were associated with IEC migration. A. Representative images for BrdU immunostaining of intestine are shown. Scale bar = 100 μm. B. Measurement of IEC migration in NEC pups. The IEC migration was defined by measuring the distance from the bottom of the crypt to the foremost labeled enterocyte (distance A) compared to the total mucosal thickness (distance B). Scale bar = 400 μm. C. IEC migration rate (defined as FLE/18). D. IEC migration (defined as FLE/total mucosal thickness 100%). E.IEC proliferation (defined as BrdU-positive cells/HPF). n= 6 animals per group, 6 fields/animal. Data represent the mean ± SEM. **P < 0.01, ****P < 0.0001 vs WT/DF; ^##^P < 0.01, ^####^P < 0.0001 vs WT/NEC; ^^P < 0.01, ^^^^P < 0.0001 vs Grx1^-/-^/NEC; one-way ANOVA.

To determine whether the IEC proliferation in Grx1 ablation pups is mediated by HIF-1α signaling, the Grx1 knockout pups were further administrated with YC-1. We found that the HIF-1α inhibition abrogated the protective effect of Grx1 ablation on IEC migration and proliferation in pups submitted to the NEC protocol (Fig. 4C, 4D, 4E).

### Stabilize HIF-1α by Grx1 ablation increases intestinal VEGF expression and improves intestinal vascular development in NEC mice, and both effects are abrogated by HIF-1 Inhibition

To explore the essential role of HIF-1α during NEC, we investigated the effects of HIF-1α on target gene VEGFA, that regulate intestinal vascular development. Both the RT-PCR and Western blot analysis methods have showed that NEC-inducing regimen significantly decreased the levels of VEGFA expression, but this trend was mitigated by Grx1 ablation (Fig. 5A, 5B), confirming the VEGFA regulated by Grx1 at the gene level. To determine whether the activation of VEGFA in Grx1 knockout pups is modulated by HIF-1α signaling, Grx1 knockout pups were administrated with the HIF-1α inhibitor, YC-1. we found that HIF-1α inhibition abrogated the activate effect of Grx1 knockout on VEGFA in pups stressed with the NEC protocol (Fig. 5A, 5B).

**Fig. 5.**
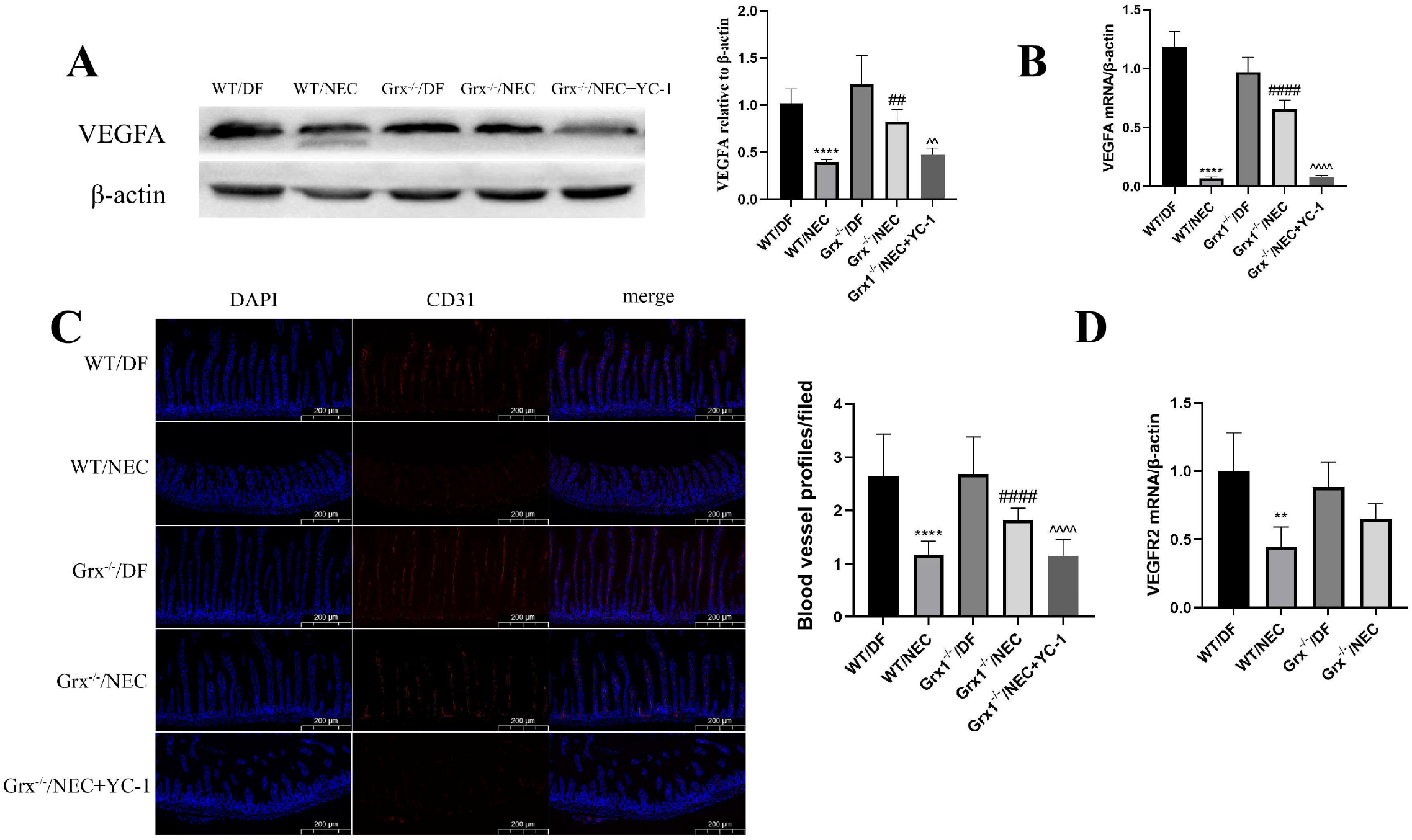
Intestinal angiogenesis in the pathogenesis of NEC was involved in VEGFR2 and VEGFA signal A. Western blot analyses for VEGFA in the pups treated as indicated. Right panel: The relative intensity of VEGFA to β-actin. B. VEGFA mRNA of intestinal tissue were detected by RT-PCR. C. Representative IF staining of CD31 for intestinal angiogenesis. Scale bar = 200 μm. Right panel: Quantitative results of panel C. D. VEGFR2 RNA was detected in the intestinal tissues by RT-PCR. Data represent the mean ± SEM. **P < 0.01, ****P < 0.0001 vs WT/DF; ^##^P < 0.01, ^###^P < 0.001, ^####^P < 0.0001 vs WT/NEC; ^^P < 0.01, ^^P < 0.001, ^^^^P < 0.0001 vs Grx1-/-/NEC; one-way ANOVA.

Previous research indicated that in experimental NEC, the intestinal microvascular development is impaired, which is regulated mainly by VEGF[21]. We next explored the involvement of HIF-1α with vessel development.

NEC stressors decreased intestinal capillary density of mice assessed by CD31 fluorescence staining, but this trend was mitigated by Grx1 ablation. However, HIF-1α inhibition by YC-1 administration abrogated the proangiogenesis effect of Grx1 ablation on NEC pups (Fig. 5C). the expression of VEGFR2 has decreased after NEC treatment, but that was not affected by Grx1 ablation (Fig. 5D).

Collectively, our results suggested that Grx1 was important in regulating intestinal capillary development in response to the NEC pathogenesis, that may be associated with the Grx1-HIF-VEGF axis.

## Discussion

To date, the potential involvement of HIF-1α in the intestinal injury of NEC has not been well defined. We here demonstrated that HIF-1α stabilization by Grx1 ablation conferred protection against the experimental NEC by reserving villi microvasculature and ameliorating intestinal injury in the immature intestine. Grx1 ablation stabilized HIF-1α by increasing GSH adducts, that promoted the expression of HIF-1α. following the HIF-1α signal, the angiogenic factor VEGFA was dependent with HIF-1α, that promoted the intestinal revascularization. The hypoxia for intestinal cells plays an essential role in development of NEC, as proved by the risk factors for NEC, like birth asphyxia, intrauterine growth restriction, congenital heart disease, and so on[1, 4]. HIF-1 present in all mammals as a highly conserved transcription factor that regulate the cellular response to ischemia and hypoxia [7]. Studies showed that HIF-1α are highly presented in ischemic bowel, and the content of serum HIF-1α was significantly upregulated in NEC patients[22, 23]. In experimental NEC, intestinal HIF-1α was significant activated[2]. But, the real role of the intestinal HIF-1α for protection or injury in murine NEC remains to be defined. Contrary to previous reports, we found that HIF-1α protein expression is decreased during NEC development in NEC pups and human NEC patients. In order to explore the reasons for the HIF-1α decline, we use immunofluorescence stain to localization HIF-1α in intestinal. We found that the HIF-1α mainly expression in intestinal villi epithelial and NEC treatment significantly decreased the intensity of fluorescence. There may be two reasons for explain this phenomenon. First, in our pathological observations, NEC is normally associated with villus damage, sloughing of the villi, which reduced the number of intestinal villi epithelial where is the site of HIF-1α mainly expression, so the total expression level of HIF-1α decreased. Second, NEC stress may have induced an early hypoxia stimulation which might account for high level HIF-1α activation and diminished subsequent over time even less than the normal level.

The glutaredoxin-1 (Grx1) is a cytosolic thioltransferase, which could efficiently catalyze GSH-protein adducts reduction[10]. Previous research in the ischemic hind limb have suggested that lack of Grx1 improved the microcirculation in the hind limb [23]. Grx1 knockout improved revascularization through GSH adducts modulation in ischemic limb to regulate HIF-1α and angiogenic genes[12], and GSH adducts modified HIF-1α stability to promote pulmonary angiogenesis[24]. Grx1-overexpressing mice in hind limb ischemia model showed impaired EC migration and attenuated revascularization[13]. These reports suggest a potential protection of Grx1 ablation to organs circulation. To explore the role of Grx1 on NEC, we used Grx1 knockout mice to establish NEC model and demonstrated that Grx1 knockout promoted revascularization and protected the intestine injury from NEC stress. Furthermore, we demonstrated that the expression of HIF-1α increased in Grx1 knockout NEC model in association with increased GSH-protein adducts. GSH-protein adducts could conserve the HIF-1α degradation and that Grx1 ablation might contribute the HIF-1α activity through preserve the GSH adducts[12]. Thus, Grx1-regulated GSH-protein adducts play a critical role in NEC pathophysiological damage. Therefore, Grx1 ablation itself could confer protection for NEC through intestinal revascularization.

The primary defensive line originate from the intestinal epithelial cells that prevent from massive pathogen invasion [25]. Human intestinal epithelium impaired can cause multiple pathophysiologic alterations, resulting in increased intestinal permeability, pathogen invasion, and intestinal damage, which is closely related to the development of NEC[26]. Since HIF-1α plays an essential role in intestinal epithelial cell physiology[27], we also explored whether HIF-1α regulated by Grx1 affecting the intestinal epithelium. We found that Grx1 ablation promoted the intestinal epithelial cell proliferation, decline in NEC pathogenesis, which is consistent with previous reports that stabilizing HIF-1α by DMOG increased intestinal epithelial cell proliferation in NEC pups[14]. That may be as the improvement of intestinal perfusion induced by Grx1 ablation promoted the intestinal epithelial cell proliferation during NEC, that sustained the epithelial integrity to protect NEC.

The chemical inhibitors of HIF-1α, 3-(5-hydroxymethy-2-furyl)-1-benzylindazole (YC-1), was confirmed to block expression of HIF-1α, so could be a potential therapeutic agent for circulation disorders[28, 29]. We demonstrated that inhibition of HIF-1α by YC-1 abolished the protection by Grx1 ablation and resulted in poor intestinal morphology in NEC pups. Furthermore, administrating YC-1 also abolished the proangiogenic effect of Grx1 ablation. This finding suggested that the beneficial effect of Grx1 ablation require HIF-1α, which is consistent with previous reports that stabilizing HIF-1α protects against NEC[14].

In response to hypoxic stress, HIF-1α regulates the downstream signal to mediate angiogenesis, including VEGF, which promotes endothelial cell proliferation and migration[30]. HIF-1α positively regulate VEGF that decreased in experimental NEC mice and in human NEC patients[31]. Consistant with previous research, the current research demonstrated that the intestinal HIF-1α expression did promote the expression of VEGF signal cascade in the condition of NEC pathological stress. In our murine NEC model, we demonstrated that VEGF, a downstream regulator of HIF-1α, are decreased during NEC development. Furthermore, Grx1 ablation improved the expression of VEGF at gene and protein level which correspond with the trend of HIF-1α. We speculated that the activation of HIF-1α induced by Grx1 ablation promoted the expression of VEGF in NEC mice.

Many studies have confirmed that HIF-1α has the positive effect on NEC, but others hold the opposite view. A study showed that dimethyloxalylglycine (DMOG), a prolyl hydroxylase (PHD) inhibitor, could stabilize HIF-1α then promote intestinal microvascular integrity through VEGF signaling[14]. Another research demonstrated that inactivation of HIF-1α by SIRT1 overexpression alleviates the inflammation associated intestinal epithelial damage[32]. Those studies show completely opposite results, may be due to different experimental methods. During angiogenesis and vasculogenesis, the VEGF modulate the activities of several kinases through its ligand, VEGFR, and ultimately guides endothelial proliferation, migration[33]. Researches shown that the microvasculature of the intestinal villi decreased in NEC pups and intestinal arterioles of NEC animals were significantly smaller compared with controls, suggesting that intestinal microcirculation is the essential component of mechanism in the development of NEC[21, 34]. These studies confirms that microvascular in injured intestine is compromised. We here indicated that the Grx1 ablation promotes villi vascular development in mice under NEC stress, which may be related to the activation of HIF-1α/VEGF signal path that induced by Grx1 ablation. Although the role of HIFs in NEC is not well addressed, accumulating evidence supported the notion that hypoxia-induced changes in HIF-mediated downstream signal factors are critical in intestinal diseases induced by ischemia and inflammation[35, 36]. Th present results suggested that the HIF/VEGF signal path may be, at least in part, modulated by Grx1 ablation to improve intestinal villus epithelial cell proliferation and microvascular integrity. In the future, further studies should investigate other mediators, which deliver the protective effects of HIF-1α.

In conclusion, GSH adducts that induced by Grx1 deletion improve intestinal revascularization via HIF/VEGF signal path in NEC mice. GSH adducts should be a potential therapeutic strategy to induce angiogenesis in intestinal to protect NEC.

## Acknowledgements

We thank Miss Siqin Yang for academic support and Jiaren Liu at the Harvard University, USA, for help with the linguistic revision of the manuscript.

## Ethics approval

All experiments were approved by the animal care and use committee of Chongqing Medical University.

